# Structural variation in *Drosophila melanogaster* spermathecal ducts and its association with sperm competition dynamics

**DOI:** 10.1101/808451

**Authors:** Ben R. Hopkins, Irem Sepil, Stuart Wigby

## Abstract

The ability of female insects to retain and use sperm for days, months, or even years after mating requires specialised storage organs in the reproductive tract. In most orders these organs include a pair of sclerotised capsules known as spermathecae. Here, we report that some *Drosophila melanogaster* females exhibit previously uncharacterised structures within the distal portion of the muscular duct that links a spermatheca to the uterus. We find that these ‘spermathecal duct presences’ (SDPs) may form in either or both ducts and can extend from the duct into the sperm-storing capsule itself. We further find that the incidence of SDPs varies significantly between genotypes, but does not change significantly with the age or mating status of females, the latter indicating that SDPs are not composed of or stimulated by sperm or male seminal proteins. We show that SDPs affect neither the number of first male sperm held in a spermatheca nor the number of offspring produced after a single mating. However, we find evidence that SDPs are associated with a lack of second male sperm in the spermathecae after females remate. This raises the possibility that SDPs provide a mechanism for variation in sperm competition outcome among females.

## 1. Introduction

Female insects commonly store sperm after mating [1]. In some species storage is particularly protracted: eusocial Hymenoptera queens can use sperm received a decade previously [2]. Where females mate multiply, sperm from rival males may be stored simultaneously and have to compete over access to limited fertilisations – a process known as sperm competition [3,4]. How the physiology of sperm storage influences the outcome of sperm competition remains a major question in the field of evolutionary reproductive biology [5,6].

Maintaining the long-term viability of sperm presents challenges [7]. Without protection, stored sperm are at risk of desiccation, thermal stress, immune attack, and the mutagenic action of oxidative stress. Retaining sperm within specialised storage organs is thought to help buffer against these effects. In most insect orders the storage organs are sclerotised capsules known as spermathecae, the number and morphology of which varies between species [8]. These organs show clear adaptations to long-term sperm use including tight control of sperm release [9] and the production of viability-enhancing secretions [10,11]. Consequently, variation in the physiology of sperm storage organs is likely to have correlated effects on female reproductive performance.

In *Drosophila melanogaster*, females store the majority of received sperm in an elongated tube known as the seminal receptacle, a novel structure found only in certain acalyptrate Dipteran families [8,12]. Genetic variation in seminal receptacle morphology has known consequences for sperm competition outcome: longer seminal receptacles increase the advantage that long sperm have over shorter rivals in both displacing sperm from storage and themselves resisting displacement [13,14]. The remaining sperm are stored in two (or rarely three) spermathecae, which consist of chitinised capsules that connect to the uterus via a muscular duct [15,16]. Sperm stored within the spermathecae can be displaced by an incoming ejaculate, although displacement here appears to be a less important contributor to paternity share compared to displacement from the seminal receptacle – at least in the short-term [5]. While variation in the morphology of *D. melanogaster* spermathecae is largely uncharacterised, there is evidence of between-population divergence in spermathecal shape in *D. affinis*, another member of the *Sophophora* subgenus to which *D. melanogaster* belongs [17]. But whether this variation has consequences for sperm storage patterns or sperm competition outcome remains untested.

Here, we report the identification of novel structures in the spermathecal duct of a subset of *D. melanogaster* females. We test for differences in the incidence of these ‘spermathecal duct presences’ (SDPs) between female genotypes, age classes, and between mated and virgin females. We then test the hypotheses that SDPs are associated with compromised sperm release, storage, and offspring production, features that would implicate them in determining sperm competition outcome.

## 2. Materials and methods

### 2.1 Fly stocks and husbandry

We used females from both wild-type Dahomey and *w*^*1118*^ backgrounds. Where females were mated, their partners were either Dahomey males or *w*^*1118*^ males expressing a *GFP-ProtB* construct [5], which fluorescently labels sperm heads green. All *GFP-ProtB* matings were with Dahomey females. For females that were double-mated, the second mating was to a Dahomey male into which *RFP-ProtB*, which labels sperm heads red [5], was previously backcrossed. All flies were reared under standardised larval densities (∼200 eggs) in bottles containing Lewis medium [18]. We collected adults as virgins under ice anaesthesia, separating them into groups of 8-12 in vials containing Lewis medium supplemented with *ad libitum* yeast granules. All flies were maintained at 25°C on a 12:12 light: dark cycle.

### 2.2 Experimental procedures

5-day old virgin females from the two genotypes (Dahomey or *w*^*1118*^) were individually isolated in yeasted vials under ice anaesthesia. Females were randomly allocated to mated or virgin treatments, one of three age classes for when they were to be dissected (1-day, 5-days, 9-days after the experimental matings), and, for the Dahomey females, whether their first (or only) male partner would transfer green fluorescent or non-fluorescent sperm.

24 hours later we aspirated males of the relevant genotype into the mating treatment vials where they remained until the pair mated. We then transferred females into fresh, yeasted vials every 24 hours for the first 3 days, and every 2 days thereafter. Additionally, on day 9 we offered 30 GFP-male-mated females the opportunity to remate with a male transferring RFP-tagged sperm. 20 mated within the 4 hours offered and were dissected 24 hours later. All female-housing vials were retained to allow any offspring to develop and were frozen once they had eclosed ready for counting. Female reproductive tracts were dissected in PBS and imaged using a Motic BA210. Fluorescent sperm counting was conducted at 40X magnification. In a second experiment, we kept virgin Dahomey females in groups of 8-12 for either 7 or 26 days, flipping them onto fresh, yeasted vials every few days, to test for later life effects.

### 2.3 Statistical analysis

All analyses were performed in R (version 3.5.1). We analysed the probability of a female exhibiting an SDP using a generalized linear model with a binary distribution, including age (as an ordered factor), female genotype, and mated status as co-factors. We used a Chi-square test to analyse differences in the proportion of 7- and 26-day old Dahomey virgin females exhibiting SDPs. When analysing sperm numbers in fluorescent-mated females, we removed 5 individuals that failed to produce any offspring (5 out of 78), which is suggestive of mating failure or infertility. We analysed the number of sperm stored by females across the two spermathecae using a linear model. We used a linear mixed effects model to analyse the number of first or second male sperm in an individual spermathecae while controlling for the identity of the female as a random effect. To analyse offspring production, we used a linear model that included male genotype (i.e. Dahomey or GFP-tagged), and the presence/absence of an SDP as factors. We analysed each female age class separately due to the bimodal distribution of offspring counts in the full dataset. In all models we tested for the significance of factors using the ‘drop1’ function with a F-test when using linear models or Chi-squared with GLMs.

## 3. Results

### 3.1 Characterisation of SDPs

Most females have clear spermathecal ducts (*e.g*. Fig. 1A). However, a subset exhibits an SDP within one or both ducts (Fig. 1B-D). SDPs appear similar in colouration to the spermathecae themselves. However, they show distinct autofluorescence at wavelength 480nm (Fig. 1B-D), suggesting compositional differences. SDPs appear to form within the duct itself rather than encircling the muscular outer wall (Fig. 1C). While there often appears to be separation from the spermathecal capsule by clear duct (Fig. 1C), SDPs occasionally continue into the capsule (Fig. 1D). In such cases, the portion of the duct that telescopes into the spermathecal capsule (the ‘introvert’, [8]) displays an altered, SDP-like fluorescence pattern (Fig. 1D). SDP size and whether it extends into the spermatheca capsule is variable between and within individuals (*e.g*. Fig. 1D).

**Figure 1.**
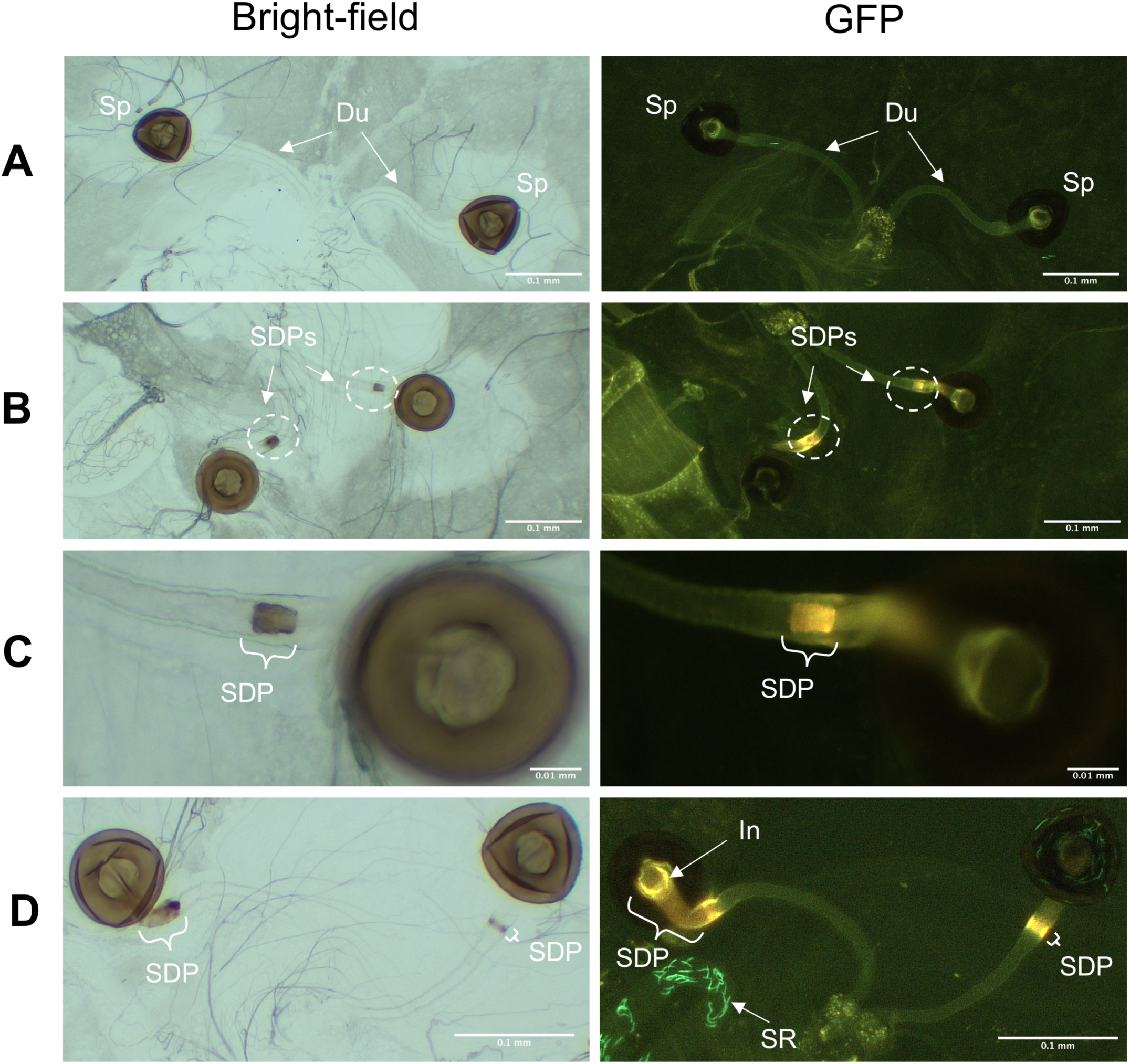
Variable spermathecal duct morphologies. (A) The spermathecae of a dissected female with clear ducts. (B) A female with spermathecal duct presences (‘SDPs’) in each duct. SDPs are circled in white. (C) The right-hand spermatheca given in (B) but at higher magnification. (D) A female showing morphologically divergent SDPs in each duct. GFP-tagged sperm are visible in some images. Sp, spermatheca; Du, spermathecal duct; SDP, spermathecal duct presence; SR, seminal receptacle; In, introvert.

### 3.2 The incidence of SDPs varies between genotypes, but is age- and mating-independent

The probability of females displaying at least one SDP was significantly higher in Dahomey compared to *w*^*1118*^ females (*LRT* =12.15, *df*=1, *p*=0.0005; Fig. 2A), but was unaffected by mating (*LRT* =0.223, *df*=1, *p*=0.637). Our data suggested a non-significant trend towards higher SDP incidence in older females, but we detected no significant interaction between age and female genotype (*LRT* =4.653, *df*=2, *p*=0.098) or the individual effect of age (*LRT* =3.34, *df*=2, *p*=0.188). Moreover, in a separate experiment we found no significant difference in the incidence of SDPs between 7- and 26-day old virgin Dahomey females (7-day = 4/31, 26-day = 7/31; χ^2^=0.815, *p*=0.367). Combining *p*-values [19] from the two independent age experiments supports the lack of a significant effect of age on SDP prevalence (*p* = 0.253).

**Figure 2.**
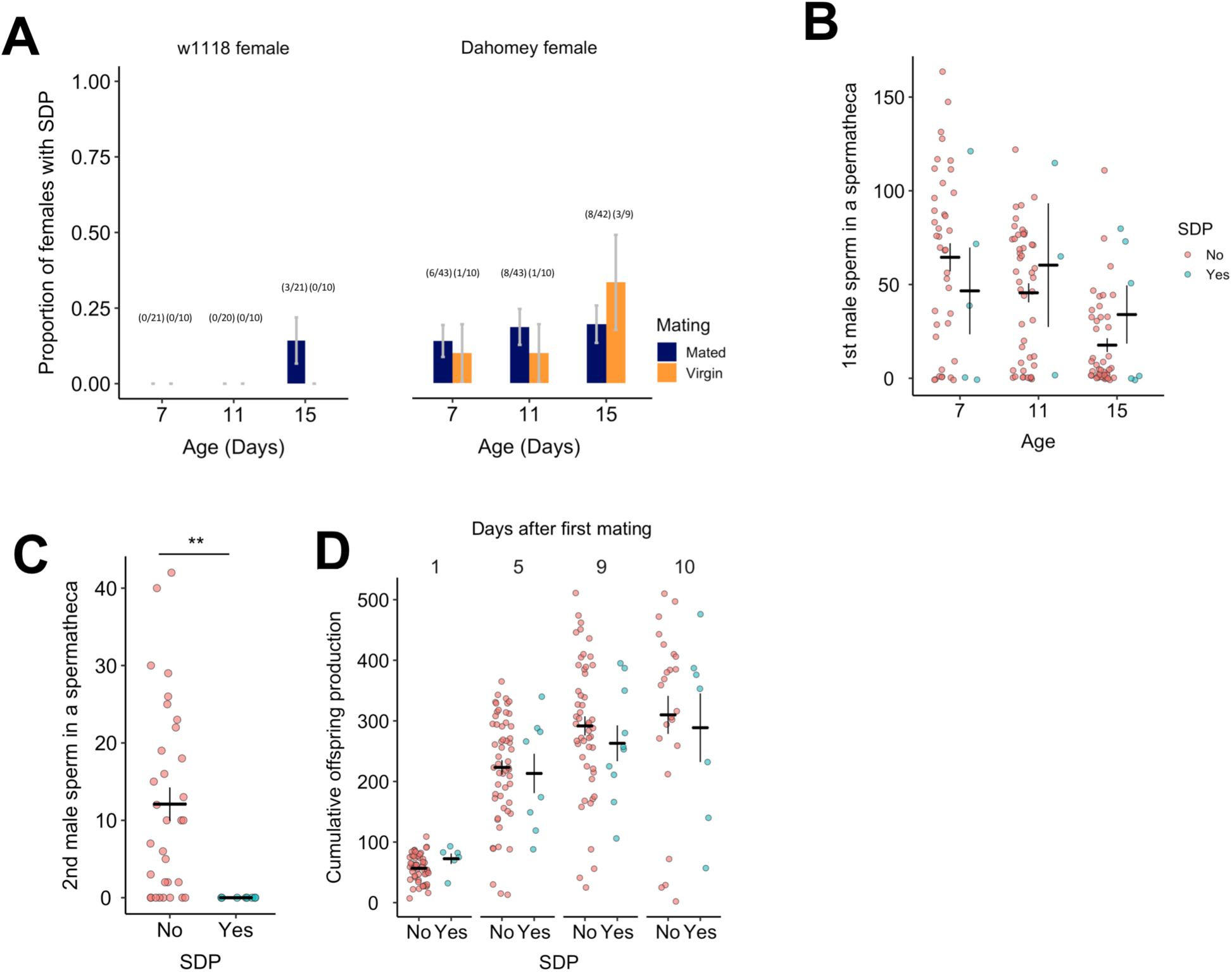
(A) The proportion of *w*^*1118*^ or Dahomey females displaying spermathecal ducts presences (SDPs) in relation to female age (in days) and mating status (mated or virgin). (B) The number of GFP-tagged sperm in a spermatheca in relation to female age and SDP presence. Females were singly mated at age 6 days. (C) The number of second male sperm in the spermathecae of females with (yes) and without (no) an SDP. (D) The cumulative number of offspring produced, plotted separately depending on whether a female had at least one SDP and in relation to the number of days after mating that the female was dissected. Both spermathecae for a single female are plotted in (B) and (C). Error bars give ±1 standard error of the sample proportion (A) or the mean (B-D). \*\**p*<0.01.

### 3.3 The presence of SDPs does not correlate with the number of sperm in the spermathecae following a single mating

The number of sperm held in individual spermathecae decreased as females aged, presumably due to use in fertilisations (F_1,69_=15.96, *p*=0.0002; Fig. 2B), but the presence of a SDP had no significant effect (F_1,122_= 0.07, *p*=0.800; Fig. 2B). These results held if we analysed the combined number of sperm held across the two spermathecae (age: F_2,68_=10.99, *p*<0.0001; SDP: F_1,68_=0.36, *p*=0.552). We found 5 cases where females produced no offspring after mating (*i.e*. were infertile or experienced mating failure), none of which exhibited an SDP. However, this did not represent a significant increase in success for females with an SDP (Fisher’s exact test, *p*=0.583).

### 3.4 SDPs inhibit second male sperm entry

There was no significant association between SDPs and whether a female remated (proportion of rematers with SDPs = 5/20; proportion of non-rematers with SDPs = 2/10; Fisher’s exact test, *p*=1). Where females remated, the number of second male sperm was significantly lower in SDP-exhibiting spermathecae (*F*_*1,31*_=8.824, *p*=0.006). All 8 of the spermathecae associated with SDPs contained 0 second male sperm. This contrasted with a range of 0 to 42 in the 32 spermathecae without SDPs. (Fig. 2C). The 8 SDP-containing spermathecae were drawn from 5 females, 3 of which exhibited SDPs in both ducts. 2 of these held no second male sperm across both spermathecae and the seminal receptacle – the only double-mating females for which this was the case.

### 3.5 SDPs do not affect the number of offspring a female produces

Combining data from all mated females, we detected no significant difference in the cumulative number of offspring produced by females with or without SDPs over any of the time points after mating (1 day: F_1,55_=0.729, *p*=0.397; 5 days: F_1,59_=1.63, *p*=0.207; 9 days: F_1,56_=1.30, *p*=0.260; 10 days: F_1,28_=0.11, *p*=0.746; Fig. 2D).

## 4. Discussion

Here, we report that the spermathecal ducts of a subset of *D. melanogaster* females contain novel structures. These ‘SDPs’ form within the distal spermathecal duct, sometimes appearing localised as a band separated from the capsule. In other cases, SDPs extend into the capsule itself. This structural variation may arise from genetic variation or represent different points in the development of the SDP: formation beginning at its most proximal location within the duct before growing upwards into the introvert and the capsule itself.

It is unclear what SDPs are made of. Superficially, they resemble the sclerotised capsule of the spermathecae, but their distinct fluorescence pattern suggests some compositional differences. Their presence in virgin females discounts a number of mechanisms through which they might form: sexually-transmitted pathogens, localised immune responses to mating [20], or via male-derived seminal products, some of which enter into the storage organs [21–23].

An important consequence of SDP formation is an apparently impaired ability for second male sperm to enter into spermathecae. While these results should be interpreted cautiously in light of the small number of individuals we found with them, they raise the possibility that SDPs affect sperm storage. Exactly why SDPs should inhibit sperm entry remains unclear. We don’t see reduced offspring production, which would be expected if there were defects in the release of resident sperm for use in fertilisations, and which might have pointed to SDPs acting as plugs. Instead, SDPs may disrupt mechanisms used to recruit sperm from the bursa, such as the release of chemoattractants or pressure changes that draw sperm into the spermathecal capsule [as in 24]. However, for this to only affect second male sperm would require either SDP formation to occur between the two matings or for SDP activity to be modified, either by the female or male-derived products, after the first mating. Given the absence of second male sperm in both the seminal receptacle and spermathecae of some SDP-bearing females, SDPs may affect sperm storage more broadly, which is conceivable given links between spermathecal activity and the viability of sperm in the seminal receptacle [25], or may themselves be a consequence of an unidentified process causing wider reproductive changes.

The between-genotype differences we detect in SDP incidence suggests that there may be standing genetic variation in populations that influences susceptibility to SDP formation. Previous work has shown that sperm competition outcome varies with female genotype [26,27] and can be subject to male x female genotype interactions [28,29]. Variation in SDP susceptibility represents a potential mechanism through which female genotype can influence sperm competition outcome. To explore this, future work should seek to identify the genetic contributions to SDP formation. Female reproductive tract genes already known to influence sperm competition outcome provide a useful starting point [26].

As females get older second male sperm precedence declines, but the underlying mechanism remains unresolved [30]. Our data show trends towards greater incidence of SDPs in older females, but any effect is small. It may be that SDP incidence is non-linear with respect to age, and accelerates much later in life than we chose to study. However, given that offspring production is concentrated in the first 3 weeks of female post-mating life (at least in the Dahomey genetic background [31]), our data covers the ages of most reproductive relevance, and it seems likely therefore that age-related changes to sperm competition outcome [e.g. 30] operate independently from SDPs.

Here, we have identified a source of structural variation in the spermathecal duct that varies among genotypes and has consequences for patterns of sperm storage. This between-female variation is likely to have important consequences for the reproductive strategies played by male *D. melanogaster*: SDPs might represent a source of variation that means second mating males cannot always enjoy the sperm competition advantage observed in *Drosophila* and many other insects [32]. Such variation might therefore allow multiple sperm competition strategies to co-circulate in populations, providing a mechanism for male x female interactions in sperm competition outcome.

## Acknowledgements

We thank Eleanor Bath, Mariana Wolfner, and Yael Heifetz for thoughtful discussion. This work was funded by the EP Abraham Cephalosporin-Oxford Graduate Scholarship to B.R.H with additional support from the BBSRC DTP. I.S. and S.W. were supported by a BBSRC fellowship to S.W. (BB/K014544/1).

## References

1. Orr TJ, Zuk M. 2012 Sperm storage. Curr. Biol. 22, R8–R10. (doi:10.1016/j.cub.2011.11.003)

2. Boomsma JJ, Baer B, Heinze J. 2005 The evolution of male traits in social insects. Annu. Rev. Entomol. 50, 395–420. (doi:10.1146/annurev.ento.50.071803.130416)

3. Parker GA. 1970 Sperm competition and its evolutionary consequences in the insects. Biol. Rev. 45, 525–567. (doi:10.1111/j.1469-185X.1970.tb01176.x)

4. Wigby S, Chapman T. 2004 Sperm competition. Curr. Biol. 14, R100–R103. (doi:10.1016/j.cub.2004.01.013)

5. Manier MK, Belote JM, Berben KS, Novikov D, Stuart WT, Pitnick S. 2010 Resolving mechanisms of competitive fertilization success in Drosophila melanogaster. Science (80-.). 328, 354–357. (doi:10.1126/science.1187096)

6. Lupold S, Pitnick S, Berben KS, Blengini CS, Belote JM, Manier MK. 2013 Female mediation of competitive fertilization success in Drosophila melanogaster. Proc. Natl. Acad. Sci. 110, 10693–10698. (doi:10.1073/pnas.1300954110)

7. Wigby S, Suarez SS, Lazzaro BP, Pizzari T, Wolfner MF. 2019 Sperm success and immunity. In Current Topics in Developmental Biology, pp. 287–313. (doi:10.1016/bs.ctdb.2019.04.002)

8. Pitnick S, Markow T, Spicer GS. 1999 Evolution of Multiple Kinds of Female Sperm-Storage Organs in Drosophila. Evolution (N. Y). 53, 1804. (doi:10.2307/2640442)

9. den Boer SPA, Baer B, Dreier S, Aron S, Nash DR, Boomsma JJ. 2009 Prudent sperm use by leaf-cutter ant queens. Proc. R. Soc. B Biol. Sci. 276, 3945–3953. (doi:10.1098/rspb.2009.1184)

10. Baer B, Eubel H, Taylor NL, O’Toole N, Millar AH. 2009 Insights into female sperm storage from the spermathecal fluid proteome of the honeybee Apis mellifera. Genome Biol. 10, R67. (doi:10.1186/gb-2009-10-6-r67)

11. den Boer SPA, Boomsma JJ, Baer B. 2009 Honey bee males and queens use glandular secretions to enhance sperm viability before and after storage. J. Insect Physiol. 55, 538–543. (doi:10.1016/j.jinsphys.2009.01.012)

12. Manier MK, Lüpold S, Pitnick S, Starmer WT. 2013 An analytical framework for estimating fertilization bias and the fertilization set from multiple sperm-storage organs. Am. Nat. 182, 552–61. (doi:10.1086/671782)

13. Miller GT, Pitnick S. 2002 Sperm-female coevolution in Drosophila. Science (80-.). 298, 1230–1233. (doi:10.1126/science.1076968)

14. Lüpold S, Manier MK, Puniamoorthy N, Schoff C, Starmer WT, Luepold SHB, Belote JM, Pitnick S. 2016 How sexual selection can drive the evolution of costly sperm ornamentation. Nature 533, 535–538. (doi:10.1038/nature18005)

15. Mayhew ML, Merritt DJ. 2013 The morphogenesis of spermathecae and spermathecal glands in Drosophila melanogaster. Arthropod Struct. Dev. (doi:10.1016/j.asd.2013.07.002)

16. Bangham J, Chapman T, Smith HK, Partridge L. 2003 Influence of female reproductive anatomy on the outcome of sperm competition in Drosophila melanogaster. Proc. R. Soc. London. Ser. B Biol. Sci. 270, 523–530. (doi:10.1098/rspb.2002.2237)

17. Miller DD. 1955 Spermatheca shape variation in Drosophila affinis. J. Hered. 46, 271–276. (doi:10.1093/oxfordjournals.jhered.a106577)

18. Clancy DJ, Kennington WJ. 2001 A simple method to achieve consistent larval density in bottle culture. Drosoph. Inf. Serv. 84, 168–169.

19. Sokal RR, Rohlf FJ. 1995 Combining probabilities from tests of significance. In Biometry: the principles and practice of statistics in biological research, pp. 794–797.

20. Carmel I, Tram U, Heifetz Y. 2016 Mating induces developmental changes in the insect female reproductive tract. Curr. Opin. Insect Sci. 13, 106–113. (doi:10.1016/j.cois.2016.03.002)

21. Avila FW, Wolfner MF. 2017 Cleavage of the Drosophila seminal protein Acp36DE in mated females enhances its sperm storage activity. J. Insect Physiol. 101, 66–72 (doi:10.1016/j.jinsphys.2017.06.015)

22. Singh A, Buehner NA, Lin H, Baranowski KJ, Findlay GD, Wolfner MF. 2018 Long-term interaction between Drosophila sperm and sex peptide is mediated by other seminal proteins that bind only transiently to sperm. Insect Biochem. Mol. Biol. 102, 43–51. (doi:10.1016/j.ibmb.2018.09.004)

23. Ram KR, Wolfner MF. 2009 A network of interactions among seminal proteins underlies the long-term postmating response in Drosophila. Proc. Natl. Acad. Sci. 106, 15384–15389. (doi:10.1073/pnas.0902923106)

24. Degner EC, Harrington LC. 2016 A mosquito sperm’s journey from male ejaculate to egg: Mechanisms, molecules, and methods for exploration. Mol. Reprod. Dev. 83, 897–911. (doi:10.1002/mrd.22653)

25. Schnakenberg SL, Matias WR, Siegal ML. 2011 Sperm-storage defects and live birth in Drosophila females lacking spermathecal secretory cells. PLoS Biol. 9, e1001192. (doi:10.1371/journal.pbio.1001192)

26. Chow CY, Wolfner MF, Clark AG. 2013 Large Neurological Component to Genetic Differences Underlying Biased Sperm Use in Drosophila. Genetics 193, 177–185. (doi:10.1534/genetics.112.146357)

27. Chen DS, Delbare SYN, White SL, Sitnik J, Chatterjee M, DoBell E, Weiss O, Clark AG, Wolfner MF. 2019 Female Genetic Contributions to Sperm Competition in *Drosophila melanogaster*. Genetics 212, 789–800. (doi:10.1534/genetics.119.302284)

28. Clark AG, Begun DJ, Prout T. 1999 Female x male interactions in Drosophila sperm competition. Science (80-.). 283, 217–220. (doi:10.1126/science.283.5399.217)

29. Chow CY, Wolfner MF, Clark AG. 2010 The genetic basis for male x female interactions underlying variation in reproductive phenotypes of Drosophila. Genetics 186, 1355–65. (doi:10.1534/genetics.110.123174)

30. Mack PD, Priest NK, Promislow DEL. 2003 Female age and sperm competition: last-male precedence declines as female age increases. Proc. R. Soc. London. Ser. B Biol. Sci. 270, 159–165. (doi:10.1098/rspb.2002.2214)

31. Sepil I, Carazo P, Perry JC, Wigby S. 2016 Insulin signalling mediates the response to male-induced harm in female Drosophila melanogaster. Sci. Rep. 6, 30205. (doi:10.1038/srep30205)

32. Laturney M, van Eijk R, Billeter J-C. 2018 Last male sperm precedence is modulated by female remating rate in Drosophila melanogaster. Evol. Lett. 2, 180–189. (doi:10.1002/evl3.50)

